# Integrated anthocyanin profiling and transcriptome analysis identify candidate genes associated with petal color variation in *Cyclamen persicum*

**DOI:** 10.64898/2026.09.23.753367

**Authors:** Demei Xia, Guoxin Yao, Xinchen Li, Yana Zhang, Ruiqi Xing, Xinyu Ma, Xinrui Han, Qianxi Hu, Zhiwei Fu, Xin Cao, Pei Chen, Shujing Xu, Guiling Liu

**Affiliations:** College of Landscape and Horticulture, Wuhu Vocational Technical University, Wuhu 241003, China; School of Urban Construction and Ecological Technology, Shanghai Institute of Technology, Shanghai 201418, China; School of Housing, Building and Planning, Universiti Sains Malaysia, Penang, Malaysia; Haidu College, Qingdao Agricultural University, Yantai 265200, China

**Keywords:** *Cyclamen persicum*, petal coloration, anthocyanin profiling, transcriptome analysis, anthocyanidin synthase, co-expression network

## Abstract

Petal coloration is a key ornamental trait in *Cyclamen persicum* Mill. and is closely associated with anthocyanin composition and relative accumulation, yet the major anthocyanins and candidate molecular factors associated with color variation remain unclear. In this study, anthocyanin profiling and transcriptome analysis were integrated to investigate three *Cyclamen* materials with distinct petal colors, BXK, FXK, and SHXK. Anthocyanin profiling showed clear separation among the three materials, indicating that petal color divergence was associated with differences in anthocyanin composition and relative abundance. Petunidin-, peonidin-, delphinidin-,malvidin-, and cyanidin-related derivatives exhibited distinct accumulation patterns. Transcriptome analysis revealed marked transcriptional divergence, with phenylpropanoid and flavonoid biosynthesis repeatedly enriched across comparisons, whereas isoflavonoid and flavone and flavonol biosynthesis were enriched in selected comparisons. Mapping differentially expressed genes onto the anthocyanin biosynthetic pathway revealed branch-specific expression patterns of structural genes, including *CHS, F3H/F3’H, F3’5’H, DFR*, and *ANS*. Notably, Cluster-26256.0, annotated as a putative anthocyanidin synthase gene, showed a progressive increase in expression from BXK to FXK and SHXK, consistent with increasing petal pigmentation. Several DFR-related transcripts also showed relatively high expression in SHXK. Co-expression analysis linked key anthocyanins, structural genes, and transcription factors, while qRT-PCR generally confirmed RNA-seq trends. These results suggest that late anthocyanin biosynthesis, particularly ANS-related transcriptional variation, may contribute to dark red petal coloration. This study provides a metabolite–transcript framework for *Cyclamen* petal coloration and identifies ANS-related genes as priority candidates for future functional studies and flower-color breeding.

## 1. Introduction

*Cyclamen persicum* Mill., commonly known as florists’ *Cyclamen*, is an important ornamental species in the genus *Cyclamen* of the family Primulaceae. It is widely cultivated as a potted flowering plant and is valued for its distinctive floral morphology and diverse flower colors. Flower color is one of the most important ornamental traits in *Cyclamen*, directly affecting its horticultural value and market preference. Among the various floral pigments, anthocyanins are major contributors to the formation of white, pink, and dark red color variation in petals. Differences in anthocyanin composition, modification, and accumulation are therefore considered a key biochemical basis underlying color diversification in *C. persicum* flowers.

Anthocyanins are a class of flavonoid pigments synthesized through the phenylpropanoid pathway and are widely distributed in flowers, fruits, leaves, and other plant tissues [1]. In flowers, anthocyanins not only determine visible pigmentation but also contribute to color intensity and hue variation through differences in hydroxylation, methylation, glycosylation, and acylation [2]. The biosynthesis of anthocyanins involves a series of well-characterized structural genes, including chalcone synthase (*CHS*), chalcone isomerase (*CHI*), flavanone 3-hydroxylase (*F3H*), flavonoid 3’-hydroxylase (*F3’H*), flavonoid 3’,5’-hydroxylase (*F3’5’H*), dihydroflavonol 4-reductase (*DFR*), and anthocyanidin synthase/leucoanthocyanidin dioxygenase (*ANS/LDOX*) [3]. In addition to core biosynthetic steps, anthocyanin modification, particularly glycosylation mediated by uridine diphosphate-glucose:flavonoid glucosyltransferases, is essential for anthocyanin stability, solubility, vacuolar accumulation, and final color expression [4]. These processes are further regulated by transcription factors, especially members of the MYB, bHLH, and WD40 families, which form MYB–bHLH–WD40 regulatory complexes controlling anthocyanin biosynthetic gene expression [5].

Substantial progress has been made in elucidating anthocyanin biosynthesis and regulation in ornamental and horticultural plants. Recent integrated metabolome and transcriptome studies in Lagerstroemia indica have shown that petal-color differentiation is often associated with coordinated changes in anthocyanin derivatives and structural-gene expression [6,7]. Similar multi-omics strategies in Brassica napus and Rhododendron simsii further linked flower-color variation with flavonoid-pathway remodeling and candidate regulatory genes [8,9]. Studies in Eustoma grandiflorum and Impatiens balsamina also indicated that anthocyanin accumulation and related gene expression jointly contribute to petal-color differences [10,11]. Previous anthocyanin profiling and transcriptome analyses of Cyclamen flowers with contrasting petal colors identified candidate genes associated with early flavonoid biosynthesis, hydroxylation, late anthocyanin formation, and glycosylation [12]. More broadly, variation in F3’H- and F3’5’H-associated branches can alter the relative formation of cyanidin- and delphinidin-derived anthocyanins, thereby contributing to flower-color differentiation [2]. However, available studies in Cyclamen have mainly examined limited color contrasts, particularly red and white flowers, whereas the anthocyanin composition, branch-specific transcriptional variation, and candidate regulatory associations underlying white, pink, and dark red petal phenotypes remain insufficiently understood. In particular, the dominant anthocyanin derivatives associated with these petal-color classes and their relationships with structural genes, glycosyltransferase genes, and transcription factors have not yet been systematically compared.

In this study, anthocyanin profiling and transcriptome sequencing were integrated to investigate anthocyanin accumulation patterns and candidate molecular factors associated with flower color variation in white, pink, and dark red *C. persicum* petals. Specifically, this study aimed to address three questions: which anthocyanin derivatives are associated with white, pink, and dark red petal phenotypes; which structural genes show branch-specific expression patterns in the anthocyanin biosynthetic pathway; and which candidate transcription factors may be associated with anthocyanin accumulation in *Cyclamen* petals. These results provide a metabolite–transcript framework for identifying candidate metabolites and genes associated with petal color variation in *C. persicum* and offer targets for further functional studies and flower-color breeding in *Cyclamen*.

## 2. Materials and Methods

### 2.1. Plant materials

The three *Cyclamen* materials were cultivated under the same greenhouse conditions to minimize environmental effects on pigment accumulation. Petals were collected at the full-bloom stage, when flowers were fully expanded and showed stable petal coloration. Samples were collected at the same time of day, immediately frozen in liquid nitrogen, and stored at −80 ℃ until analysis.

### 2.2. RNA extraction, library construction, and sequencing

RNA sequencing was performed by Metware Biotechnology Co., Ltd. (Wuhan, China). Total RNA was extracted from petals of BXK, FXK, and SHXK using the RNAprep Pure Plant Plus Kit (DP441, TIANGEN, China) according to the manufacturer’s instructions. RNA quality and integrity were evaluated prior to library construction. After mRNA enrichment, cDNA libraries were prepared using the NEBNext Ultra RNA Library Prep Kit for Illumina (NEB, USA). Library quality was initially assessed using Qubit for concentration determination, and insert size was further examined using a fragment analyzer. Qualified libraries were then subjected to high-throughput paired-end sequencing.

### 2.3. Transcriptome assembly, expression quantification, and differential expression analysis

Raw sequencing reads were processed by quality control to obtain clean reads. Reads containing adapter sequences were removed. Paired reads were also discarded if the proportion of ambiguous nucleotides (N) exceeded 10% of the total bases in a read or if the proportion of low-quality bases (Q ≤ 20) exceeded 50% of the total bases in a read. The high-quality clean reads were de novo assembled using Trinity to generate transcript sequences, and redundant sequences were further clustered using Corset to obtain unigenes.

Gene expression levels were normalized as fragments per kilobase of transcript per million mapped reads (FPKM). Pairwise differential expression analysis among BXK, FXK, and SHXK was performed using DEGSeq. Transcripts with an absolute log2 fold change (|log2FC|) ≥ 1 and a P-value < 0.05 were defined as differentially expressed genes (DEGs).

### 2.4. KEGG pathway enrichment analysis

To investigate the biological functions of the identified DEGs, Kyoto Encyclopedia of Genes and Genomes (KEGG) pathway enrichment analysis was performed using the KEGG database and KOBAS software. Pathways with a Q-value < 0.05 were considered significantly enriched and were used to identify biological processes potentially associated with anthocyanin biosynthesis and petal coloration.

### 2.5. Anthocyanin extraction and LC–MS/MS analysis

Petal samples were ground into powder under liquid nitrogen, and 50 mg of each sample was extracted with 50% methanol containing 0.1% hydrochloric acid. The mixture was vortexed for 5 min, sonicated for 5 min, and centrifuged at 12,000 rpm for 3 min at 4 ℃. The supernatant was collected, and the extraction was repeated once under the same conditions. The two supernatants were combined, filtered through a 0.22 μm membrane, and transferred to sample vials for LC–MS/MS analysis.

Anthocyanin profiling was performed using an ultra-performance liquid chromatography–tandem mass spectrometry (UPLC–MS/MS) system consisting of an ExionLC AD UPLC system and a QTRAP 6500+ mass spectrometer [13].

Chromatographic separation was carried out on an ACQUITY BEH C18 column (1.7 μm, 2.1 mm × 100 mm). The mobile phase consisted of ultrapure water containing 0.5% formic acid (phase A) and methanol containing 0.5% formic acid (phase B). The gradient elution program was as follows: 5% B at 0.00 min, increased to 50% at 6.00 min, increased to 95% at 12.00 min, maintained at 95% for 2 min, then reduced to 5% at 14.00 min and equilibrated for 2 min. The flow rate was 0.35 mL min^-1^, the column temperature was maintained at 40 ℃, and the injection volume was 2 μL.

Mass spectrometric detection was performed in positive electrospray ionization mode. The ion source temperature was set at 550 ℃, the ion spray voltage was 5500 V, and the curtain gas was set at 35 psi. Data acquisition was conducted in multiple reaction monitoring (MRM) mode, and each ion pair was optimized based on the corresponding declustering potential and collision energy.

### 2.6. Anthocyanin identification and quantitative analysis

Anthocyanins were identified based on the MWDB database established by Metware Biotechnology Co., Ltd. (Wuhan, China) in combination with tandem mass spectrometry data. Quantitative analysis was performed in MRM mode by integrating chromatographic peak areas, with standard substances used as internal references for calibration. Hierarchical clustering analysis was conducted based on anthocyanin accumulation levels across samples. Differential anthocyanins among the three flower-color groups were screened using orthogonal partial least squares discriminant analysis (OPLS-DA).

### 2.7. qRT-PCR validation

To validate the reliability of the RNA-seq data, quantitative real-time PCR (qRT-PCR) was performed for eight selected transcripts. Total RNA was extracted from petals of BXK, FXK, and SHXK using the same RNA samples employed for transcriptome sequencing. Total RNA (1 μg) was used to synthesize first-strand cDNA using the PrimeScript™ RT Master Mix kit (TaKaRa, Dalian, China) according to the manufacturer’s instructions. The qRT-PCR assays were performed using 2 × ChamQ Universal SYBR qPCR Master Mix (Vazyme, Nanjing, China) on a C1000 Touch™ Thermal Cycler system (Bio-Rad, Hercules, CA, USA). The gene-specific primers used in this study are listed in Table S1. The eEF1*α* gene was used as the internal reference gene for normalization. Relative transcript levels were calculated using the 2^−ΔΔCt^ method, and primer specificity was confirmed by the presence of a single peak in the melting-curve analysis. Three biological replicates were analyzed for each material.

### 2.8. Co-expression network construction and statistical analysis

An integrated co-expression network was constructed to investigate the regulatory relationships among differentially accumulated anthocyanins, structural genes involved in anthocyanin biosynthesis, and differentially expressed transcription factors [14]. Pearson’s correlation coefficients were calculated among these variables, and significantly correlated pairs were retained for network construction. The resulting network was visualized using Cytoscape.

All experiments were conducted with three biological replicates, and data are presented as the mean ± standard deviation (SD). The statistical thresholds for differential gene expression analysis, KEGG enrichment analysis, and differential anthocyanin screening are described in the corresponding sections above.

## 3. Results

### 3.1. Anthocyanin profiling reveals distinct accumulation patterns in Cyclamen petals

Clear phenotypic differences were observed among BXK, FXK, and SHXK, with BXK exhibiting white petals, FXK showing pink petals, and SHXK displaying dark red petals (Fig. 1A). To investigate the metabolic basis underlying these color differences, anthocyanin profiling was performed and the data were subjected to orthogonal partial least squares discriminant analysis (OPLS-DA). A total of 61 anthocyanin-related metabolites were detected in the three *Cyclamen* materials, including 14 cyanidin-related, 12 peonidin-related, 9 delphinidin-related, 8 pelargonidin-related, 7 petunidin-related, 6 malvidin-related, 3 procyanidin-related, and 2 other flavonoid compounds. The three groups were clearly separated, and the biological replicates clustered tightly within each group, indicating marked differences in anthocyanin composition and good reproducibility of the metabolomic data (Fig. 1B).

**Fig. 1.**
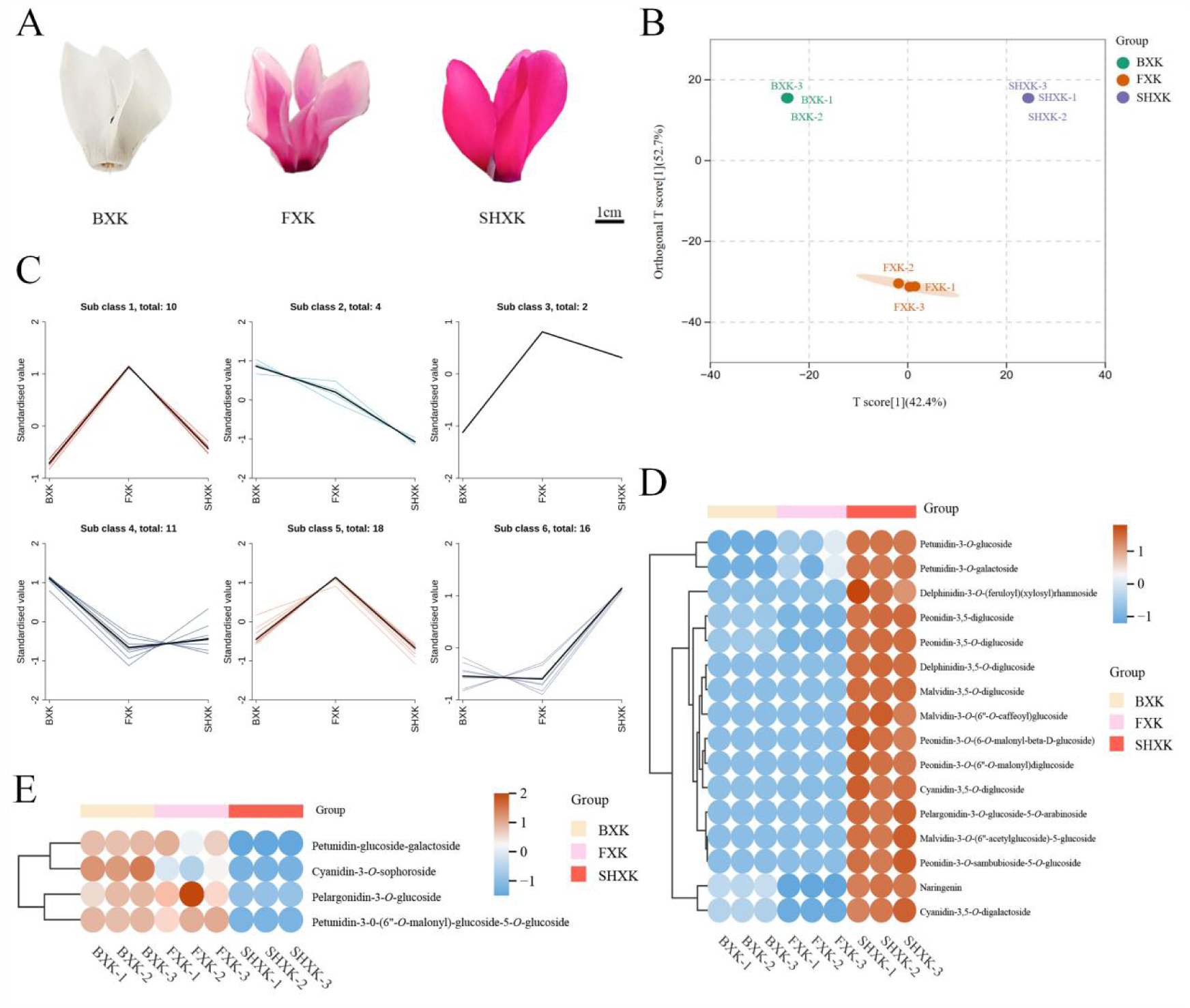
Anthocyanin profiling reveals distinct accumulation patterns in *Cyclamen* petals. (A) Petal phenotypes of the three *Cyclamen* materials, BXK, FXK, and SHXK. Scale bar = 1 cm. (B) Orthogonal partial least squares discriminant analysis (OPLS-DA) score plot based on anthocyanin profiling data, showing clear separation among BXK, FXK, and SHXK. (C) K-means clustering analysis of all detected anthocyanin-related metabolites. Six subclasses were identified according to their standardized accumulation patterns among the three materials.(D) Heatmap showing anthocyanins predominantly accumulated in SHXK.(E) Heatmap showing anthocyanins with relatively higher accumulation in BXK and/or FXK than in SHXK. In the heatmaps, columns represent biological replicates and rows represent anthocyanin metabolites. Color scales indicate relative accumulation levels.

Based on their standardized accumulation patterns across the three materials, all 61 anthocyanin-related metabolites were classified into six subclasses by K-means clustering (Fig. 1C). Subclasses 1 and 5 showed peak accumulation in FXK, whereas subclass 6 exhibited the highest accumulation in SHXK. In contrast, subclass 2 displayed a progressive decline from BXK to SHXK, while subclass 4 showed relatively high abundance in BXK, followed by a marked decrease in FXK and a partial recovery in SHXK.

Based on the criteria of VIP ≥ 1 and FDR < 0.05, 30 anthocyanin-related metabolites were identified as significantly differential among BXK, FXK, and SHXK. Among these differential metabolites, 16 showed the highest accumulation in SHXK, 12 in BXK, and 2 in FXK, indicating that anthocyanin accumulation was differentially remodeled among the three materials.

Heatmap analysis further showed that several glycosylated anthocyanins accumulated predominantly in SHXK, including malvidin-3,5-*O*-diglucoside, peonidin-3,5-*O*-diglucoside, cyanidin-3,5-*O*-diglucoside, delphinidin-3,5-*O*-diglucoside, and malvidin-3-*O*-(6”-*O*-caffeoyl)glucoside (Fig. 1D). In particular, malvidin-3,5-*O*-diglucoside showed markedly higher relative abundance in SHXK, with an average value of 13909.72, compared with 36.58 in BXK and 41.22 in FXK. Peonidin-3,5-*O*-diglucoside, cyanidin-3,5-*O*-diglucoside, and delphinidin-3,5-*O*-diglucoside also accumulated at higher levels in SHXK, with average values of 338.63, 196.57, and 77.02, respectively. Collectively, these results indicate that the dark red phenotype of SHXK was closely associated with the preferential accumulation of multiple glycosylated anthocyanins, especially malvidin-, peonidin-, cyanidin-, and delphinidin-derived diglucosides.

### 3.2. Transcriptome profiling reveals distinct transcriptional differences in Cyclamen petals

To further explore the molecular basis of petal color variation, transcriptome sequencing was performed using BXK, FXK, and SHXK petals, and 25,109 expressed genes or transcripts were identified. Principal component analysis (PCA) showed that the biological replicates of each group clustered closely together and were clearly separated from the other groups (Fig. 2A), indicating high reproducibility of the transcriptomic data and distinct global transcriptional profiles among BXK, FXK, and SHXK. PC1 and PC2 explained 31.19% and 24.07% of the total variance, respectively, suggesting that a substantial proportion of transcriptional variation was associated with petal color differentiation.

**Fig. 2.**
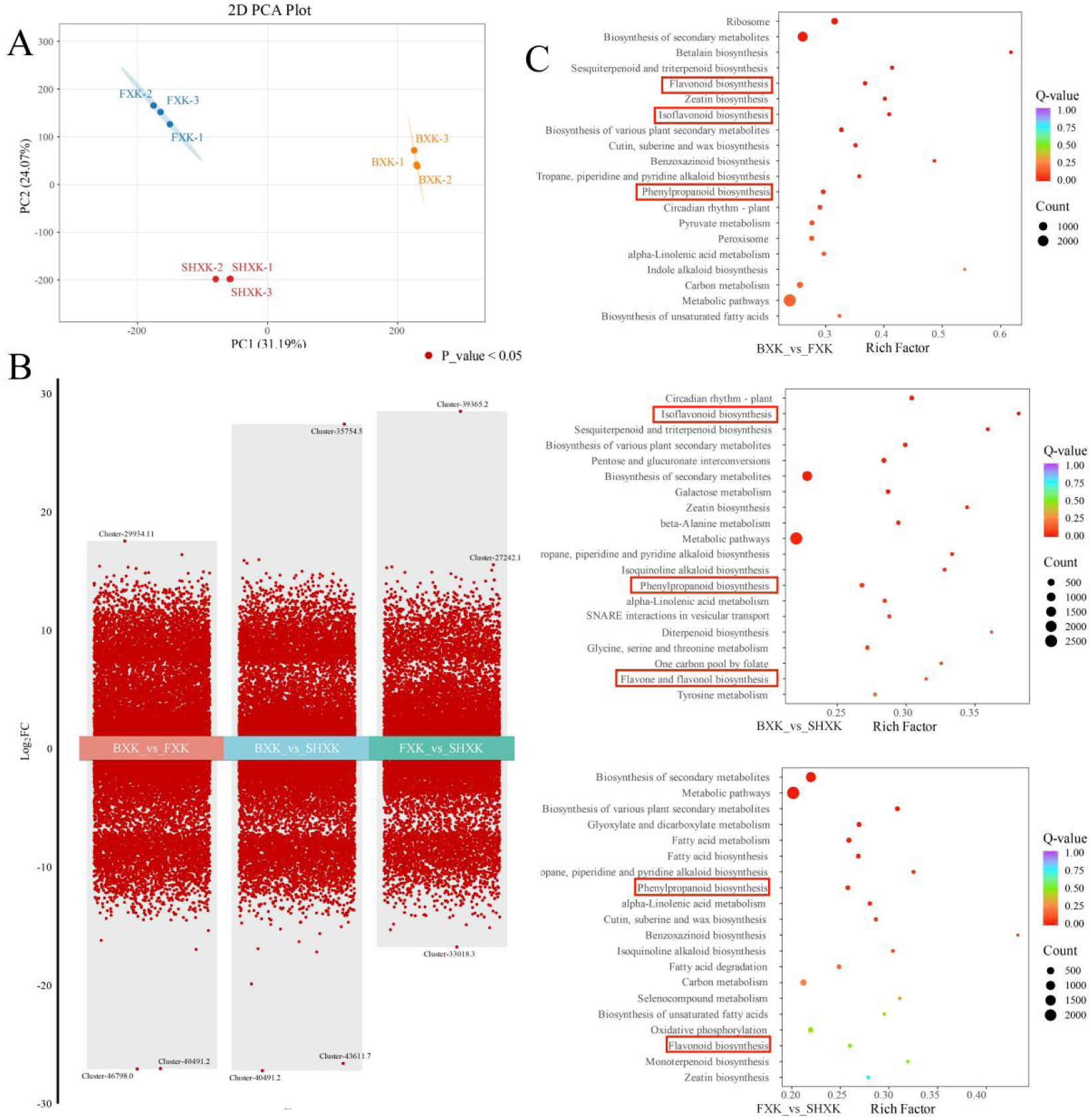
Transcriptome profiling and KEGG enrichment analysis of *Cyclamen* petals. (A) Principal component analysis (PCA) of transcriptomic data from BXK, FXK, and SHXK, showing clear separation among the three materials and good clustering of biological replicates. (B) Volcano plots showing differentially expressed genes in the BXK vs FXK, BXK vs SHXK, and FXK vs SHXK comparisons. Red dots indicate DEGs meeting the criteria of |log2FC| ≥ 1 and P-value < 0.05. (C) KEGG enrichment analysis of DEGs in the BXK vs FXK, BXK vs SHXK, and FXK vs SHXK comparisons, shown from top to bottom. Bubble size represents the number of enriched genes, and bubble color indicates the Q-value. Pathways with a Q-value < 0.05 were considered significantly enriched.

Differential expression analysis revealed extensive transcriptional differences among BXK, FXK, and SHXK (Fig. 2B). In the BXK vs FXK comparison, 17,845 DEGs were identified, including 8,288 up-regulated and 9,557 down-regulated genes. In the BXK vs SHXK comparison, 16,854 DEGs were identified, including 8,330 up-regulated and 8,524 down-regulated genes. In the FXK vs SHXK comparison, 13,969 DEGs were identified, including 7,391 up-regulated and 6,578 down-regulated genes. The widespread distribution of DEGs across the three pairwise comparisons indicated substantial transcriptional variation among the three flower-color materials.

### 3.3. KEGG enrichment analysis highlights flavonoid- and phenylpropanoid-related pathways associated with Cyclamen petal coloration

To identify the major biological pathways associated with petal color differentiation, KEGG enrichment analysis was performed for the DEGs from each pairwise comparison (Fig. 2C). The enriched pathways were mainly related to secondary metabolism, including flavonoid biosynthesis, phenylpropanoid biosynthesis, isoflavonoid biosynthesis, and flavone and flavonol biosynthesis.A total of 139 genes were annotated in flavonoid biosynthesis, 157 in phenylpropanoid biosynthesis, 59 in isoflavonoid biosynthesis, 37 in flavone and flavonol biosynthesis, and 2 in anthocyanin biosynthesis.

In the BXK vs FXK comparison, flavonoid biosynthesis, phenylpropanoid biosynthesis, isoflavonoid biosynthesis, and flavone and flavonol biosynthesis contained 114, 116, 45, and 28 DEGs, respectively. In the BXK vs SHXK comparison, these pathways contained 86, 105, 42, and 28 DEGs, respectively. In the FXK vs SHXK comparison, phenylpropanoid biosynthesis and flavonoid biosynthesis remained significantly enriched. The repeated enrichment of these pathways indicated that petal color differentiation among BXK, FXK, and SHXK was closely associated with transcriptional variation in the phenylpropanoid and flavonoid metabolic network.

Taken together, these results suggest that phenylpropanoid- and flavonoid-related pathways were closely associated with the distinct petal color phenotypes of BXK, FXK, and SHXK.

### 3.4. Differential expression of structural genes involved in the anthocyanin biosynthetic pathway in Cyclamen petals

To further clarify the transcriptional basis underlying petal color divergence, the DEGs were mapped onto the anthocyanin biosynthetic pathway (Fig. 3). Multiple CHS-related transcripts showed differential expression among BXK, FXK, and SHXK, and several showed relatively higher expression in FXK, indicating that early flavonoid biosynthetic genes exhibited distinct transcriptional profiles among the three flower-color materials. Genes involved in the downstream hydroxylation steps also displayed distinct expression patterns. These included *F3H/F3’H*-related transcripts (Cluster-44802.0, Cluster-44802.1, Cluster-44802.2, Cluster-44802.4, and Cluster-44802.5) as well as the *F3’5’H*-related transcript Cluster-27057.0. The differential expression of these genes suggested that the conversion of dihydrokaempferol into dihydroquercetin and dihydromyricetin differed among the three materials, thereby potentially affecting the relative formation of cyanidin- and delphinidin-derived anthocyanins.

**Fig. 3.**
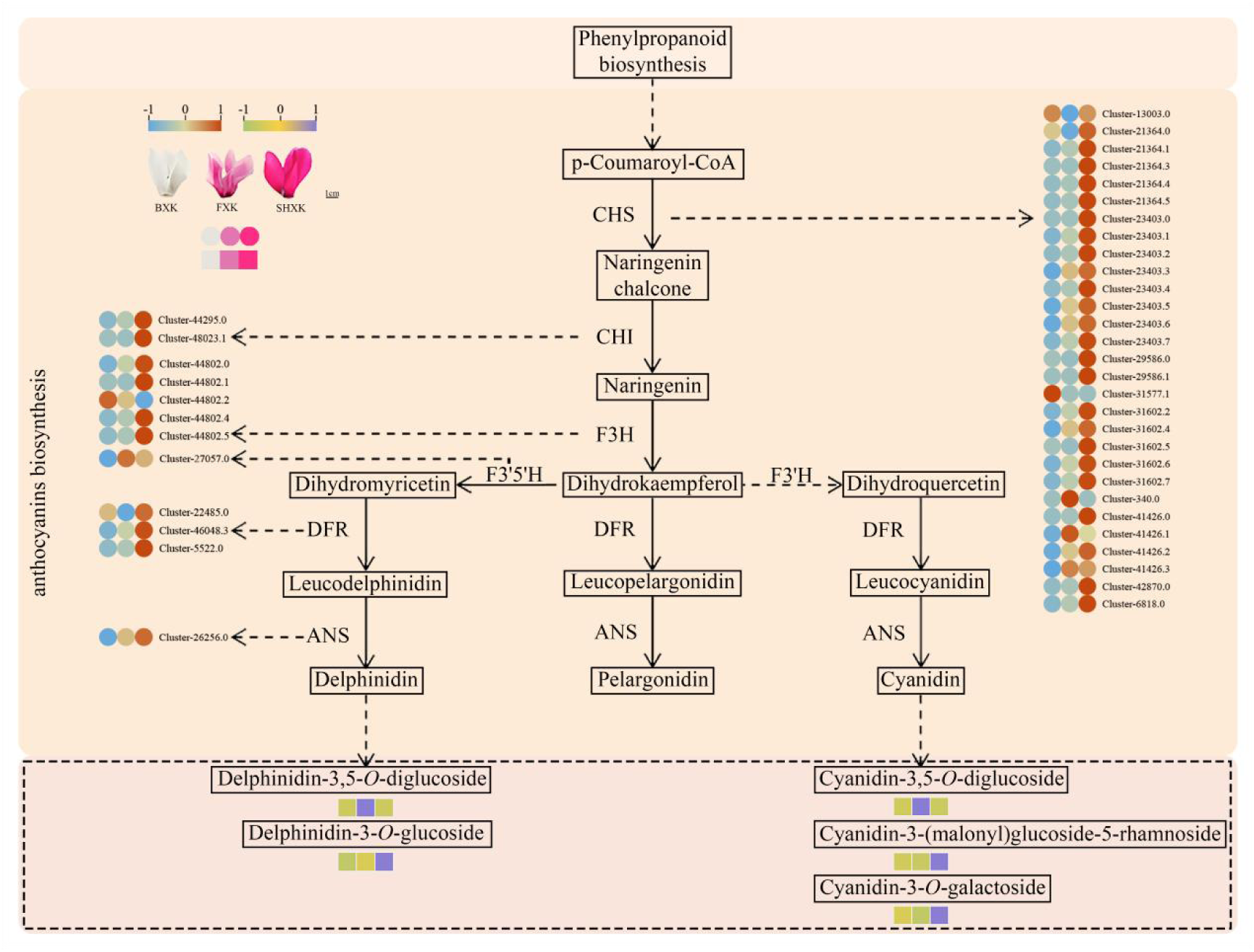
Differential expression of structural genes in the anthocyanin biosynthetic pathway of *Cyclamen*petals. Differentially expressed genes identified from transcriptome analysis were mapped onto the phenylpropanoid and anthocyanin biosynthetic pathways. Structural genes involved in early flavonoid biosynthesis, hydroxylation, and late anthocyanidin formation showed distinct expression patterns among BXK, FXK, and SHXK. Representative terminal anthocyanin derivatives are shown at the bottom of the pathway. Colored circles indicate relative gene expression levels, and colored squares indicate relative metabolite accumulation levels in BXK, FXK, and SHXK.

Further downstream, DFR-related transcripts and the ANS-related gene Cluster-26256.0 also showed expression divergence among BXK, FXK, and SHXK. Cluster-26256.0 showed relatively low expression in BXK, intermediate expression in FXK, and the highest expression in SHXK, a pattern directionally consistent with the observed white, pink, and dark red petal-color classes. In addition, the DFR-related transcripts Cluster-22485.0, Cluster-46048.3, and Cluster-5522.0 showed relatively high expression in SHXK. These expression patterns were associated with the preferential accumulation of multiple terminal anthocyanin derivatives in the dark red petals.

Together, these results indicate that petal color variation among the three materials was associated with branch-specific transcriptional differences in early flavonoid biosynthesis, hydroxylation-related genes, and late anthocyanin biosynthetic genes. In particular, Cluster-26256.0 and several DFR-related transcripts showed expression trends directionally consistent with the darker petal phenotype of SHXK.

### 3.5. Co-expression network analysis identifies candidate regulators associated with petal coloration

To further investigate the correlation-based associations underlying petal coloration in *Cyclamen*, an integrated co-expression network was constructed based on differentially accumulated anthocyanins, anthocyanin biosynthetic genes, and differentially expressed transcription factors (Fig. 4). The resulting network revealed extensive correlation-based associations among anthocyanin metabolites, biosynthetic genes, and multiple transcription factor families, suggesting that petal color variation in *Cyclamen* may be associated with coordinated expression patterns among these components.

**Fig. 4.**
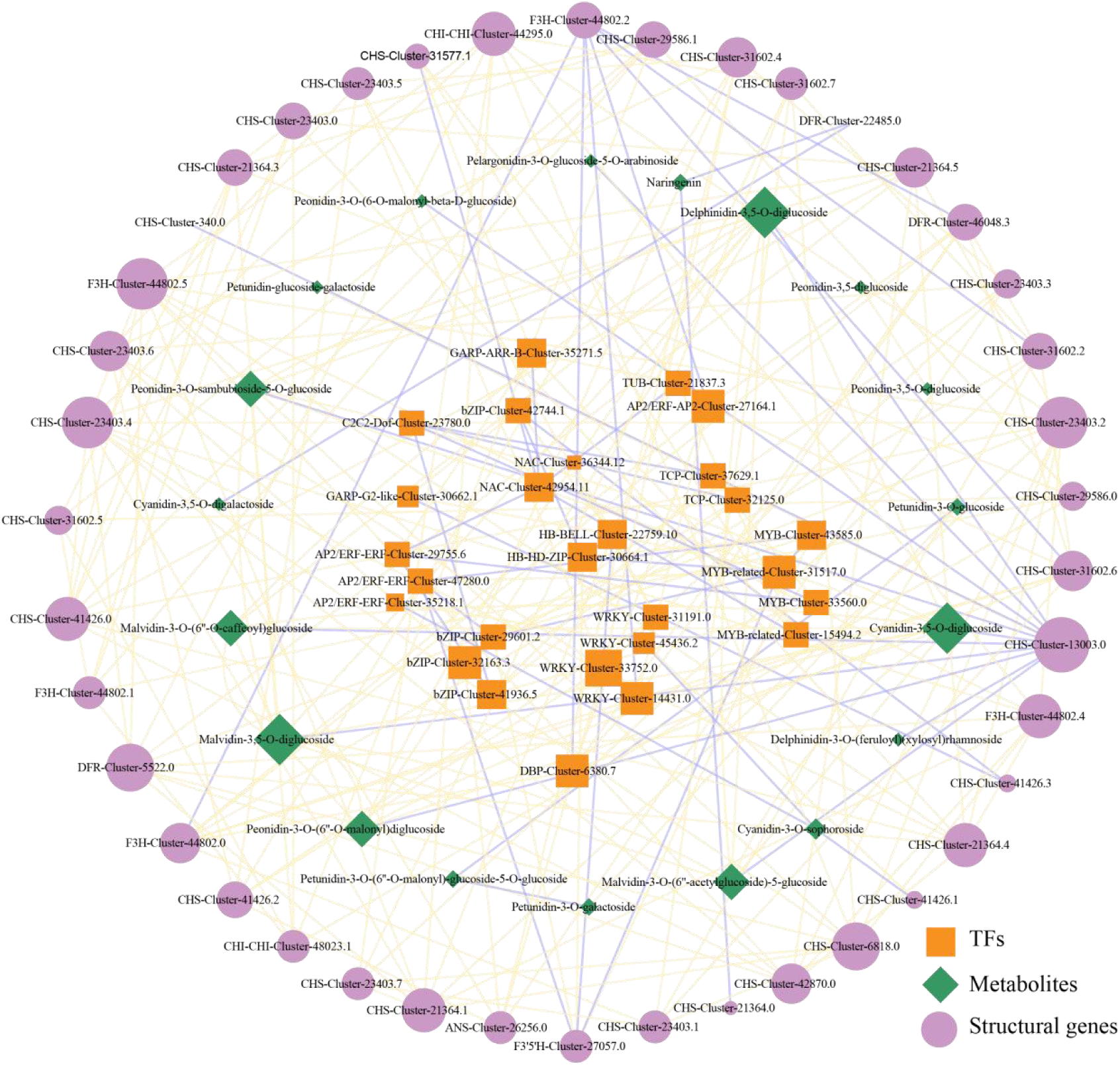
Integrated co-expression network linking anthocyanin metabolites, structural genes, and transcription factors in *Cyclamen*. The network diagram, visualized using Cytoscape, illustrates correlation-based associations among anthocyanin metabolites, structural genes, and candidate transcription factors in *Cyclamen*. The nodes represent different biological components involved in the co-expression network: Green diamonds represent key anthocyanin metabolites associated with petal coloration; purple circles represent structural genes involved in the anthocyanin biosynthetic pathway; and orange squares represent candidate transcription factors associated with anthocyanin accumulation. The edges connecting the nodes represent correlation pairs retained for network construction.

Several key anthocyanin derivatives, including delphinidin-3,5-*O*-diglucoside, cyanidin-3,5-*O*-diglucoside, malvidin-3,5-*O*-diglucoside, malvidin-3-*O*-(6’’-*O*-caffeoyl)glucoside, peonidin-3,5-*O*-diglucoside, petunidin-3-*O*-glucoside, and petunidin-3-*O*-galactoside, were closely connected with multiple structural genes and transcription factors in the network. At the biosynthetic gene level, numerous *CHS* members, together with *F3H/F3’H*-, *DFR*-, and *ANS*-related genes, were linked to these anthocyanin metabolites, further supporting their potential involvement in branch-specific anthocyanin accumulation in *Cyclamen petals*.

Notably, several MYB and MYB-related transcription factors showed relatively high connectivity with anthocyanin metabolites and structural genes, suggesting that they may represent important candidate regulators of anthocyanin accumulation. Among these candidates, MYB-Cluster-43585.0 and MYB-Cluster-33560.0 were selected as priority MYB candidates because of their expression differences among BXK, FXK, and SHXK. MYB-Cluster-43585.0 showed average FPKM values of 74.15, 161.25, and 197.11 in BXK, FXK, and SHXK, respectively, and was significantly up-regulated in both FXK vs BXK and SHXK vs BXK. Among them, several MYB- and MYB-related nodes, such as MYB-Cluster-43585.0, MYB-Cluster-33560.0, MYB-related-Cluster-31517.0, and MYB-related-Cluster-15494.2, showed particularly dense associations within the network, suggesting that they may represent priority candidate regulators associated with anthocyanin accumulation. These expression patterns suggest that MYB-Cluster-43585.0 and MYB-Cluster-33560.0 may be important candidate regulators associated with anthocyanin accumulation in colored *Cyclamen* petals.

Overall, the integrated co-expression network revealed correlation-based associations among anthocyanin metabolites, structural genes, and candidate transcription factors. These results provide a framework for prioritizing candidate regulators associated with anthocyanin accumulation for future functional validation in *Cyclamen*.

### 3.6. qRT-PCR validation of selected transcripts

To evaluate the reliability of the RNA-seq expression trends, eight candidate genes were selected for qRT-PCR analysis using eEF1*α* as the internal reference gene (Fig. 5). As shown in Fig. 5, the expression patterns determined by qRT-PCR were generally consistent with the FPKM trends obtained from RNA-seq, indicating the reliability of the transcriptomic dataset.

**Fig. 5.**
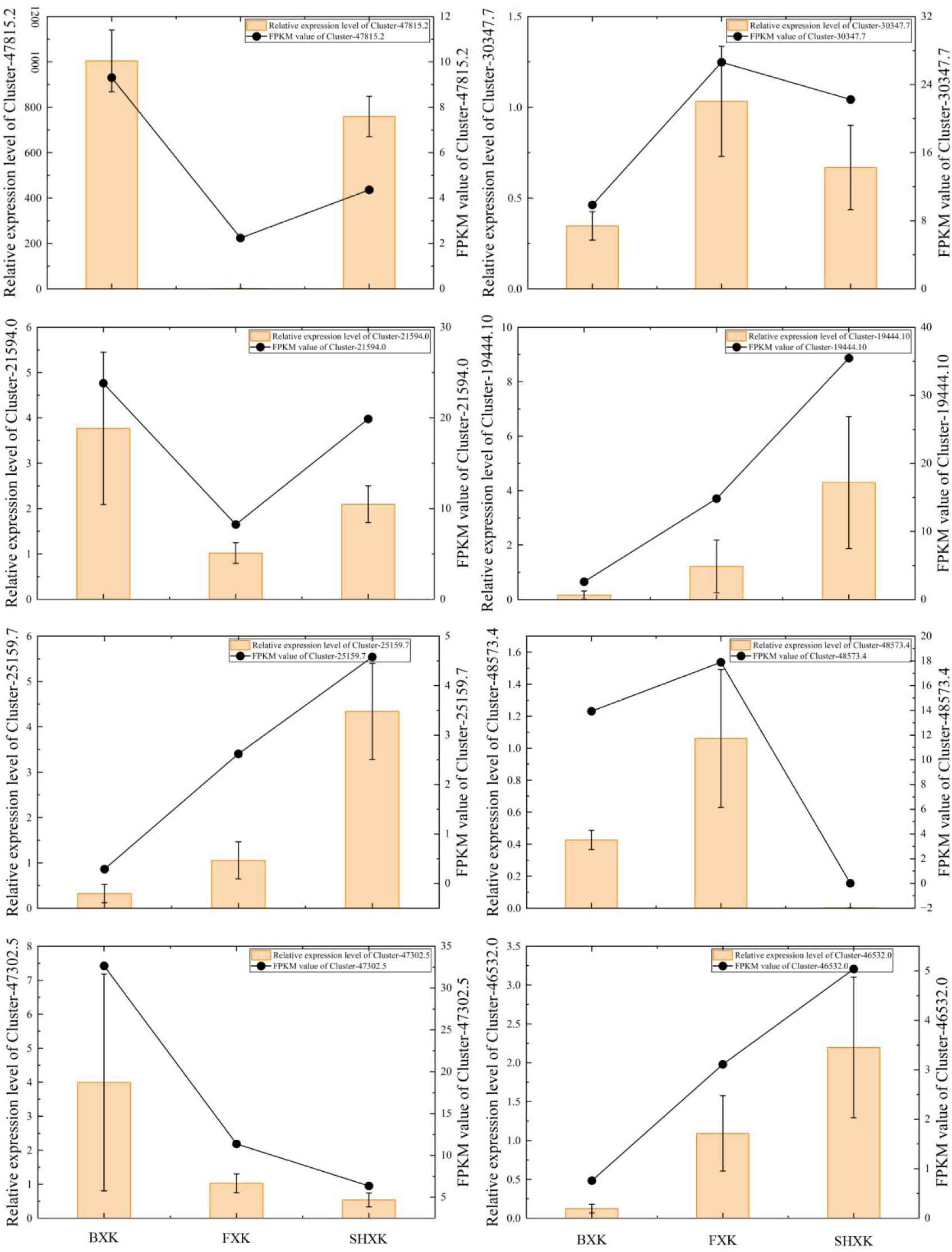
qRT-PCR validation of selected transcripts in *Cyclamen* petals. Relative expression levels of eight selected candidate genes, including Cluster-47815.2, Cluster-30347.7, Cluster-21594.0, Cluster-19444.10, Cluster-25159.7, Cluster-48573.4, Cluster-47302.5, and Cluster-46532.0, were analyzed by qRT-PCR in BXK, FXK, and SHXK. Bars represent relative expression levels determined by qRT-PCR, and error bars indicate the standard deviation of three biological replicates. The black line indicates the corresponding FPKM values obtained from RNA-seq. Overall, the qRT-PCR results were generally consistent with the transcriptome data, supporting the reliability of the RNA-seq analysis.

Among the selected genes, Cluster-30347.7 showed the highest expression level in FXK, whereas Cluster-19444.10, Cluster-25159.7, and Cluster-46532.0 exhibited progressively increased expression from BXK to SHXK, with the highest transcript abundance detected in SHXK. In contrast, Cluster-21594.0 and Cluster-47302.5 displayed relatively high expression in BXK and lower expression in FXK and SHXK.

Cluster-47815.2 also showed a higher expression level in BXK and SHXK than in FXK. In addition, Cluster-48573.4 exhibited the highest relative expression in FXK, whereas its expression was comparatively lower in BXK and SHXK. Overall, although slight differences in expression magnitude were observed between qRT-PCR and RNA-seq, the overall expression tendencies were highly consistent between the two methods.

These results generally confirmed the RNA-seq expression trends of the eight selected transcripts and supported the reliability of the transcriptomic dataset.

## 4. Discussion

### 4.1. Anthocyanin Profiles Associated with White-to-Dark Red Petal Coloration

Petal coloration in *Cyclamen* appears to be associated not only with the total accumulation of anthocyanins, but also with differences in anthocyanidin backbone composition and subsequent modification patterns. In this study, SHXK, the dark red material, showed preferential relative accumulation of several petunidin-, peonidin-, delphinidin-, malvidin-, and cyanidin-related anthocyanins. This pattern indicates that the dark red phenotype of SHXK was associated with variation across multiple anthocyanin branches rather than with the relative accumulation of a single dominant pigment.

Comparable integrated analyses in Prunus mume and Scutellaria baicalensis have shown that petal-color variation is frequently associated with changes in multiple anthocyanin derivatives rather than a single pigment class [15,16].Studies in Orychophragmus violaceus and dark-flowered ornamental crops further suggest that darker or more saturated floral phenotypes may involve coordinated variation in multiple anthocyanin branches [17,18].Similar findings in Rosa rugosa and Camellia oleifera also support the view that flower-color differences are associated with combined changes in anthocyanin composition and modification patterns [19,20].By contrast, delphinidin-, petunidin-, and malvidin-related derivatives were detected in the present Cyclamen materials, indicating that products of multiple hydroxylation and modification branches were retained in the petal anthocyanin profile.

### 4.2. Branch-specific transcriptional regulation of anthocyanin biosynthesis in Cyclamen petals

Late anthocyanin biosynthesis represents a key stage linking dihydroflavonol metabolism with the formation of colored anthocyanidins. In this pathway, DFR catalyzes the reduction of dihydroflavonols to the corresponding leucoanthocyanidins, whereas ANS subsequently converts these colorless intermediates into anthocyanidins. The coordinated expression of *DFR*- and *ANS*-related genes may therefore influence the supply of anthocyanidin precursors available for subsequent modification and accumulation in petals.

In the present study, the putative *ANS* gene Cluster-26256.0 showed relatively low expression in BXK, intermediate expression in FXK, and the highest expression in SHXK. This expression pattern was directionally consistent with the observed white, pink, and dark red petal-color classes. In addition, several *DFR*-related transcripts, particularly Cluster-46048.3 and Cluster-5522.0, showed relatively high expression in SHXK. These results indicate that transcriptional variation in late anthocyanin biosynthetic genes was associated with petal color variation among the three materials.

The metabolomic data further showed that SHXK preferentially accumulated multiple terminal anthocyanin derivatives, including malvidin-, peonidin-, cyanidin-, and delphinidin-related glycosides. Similar associations between late anthocyanin-biosynthetic genes and anthocyanin accumulation have been reported in Allium wallichii and Impatiens balsamina [21,22]. In the present study, the relative accumulation patterns of these glycosylated derivatives were directionally consistent with the expression differences observed for several DFR- and ANS-related transcripts. However, the present data do not distinguish the relative contributions of anthocyanidin formation, methylation, glycosylation, or other downstream modification steps to the accumulation of individual anthocyanin derivatives.

However, the present data do not establish a direct causal relationship between the identified transcripts and individual anthocyanin metabolites. Although several glycosylated anthocyanins accumulated in SHXK, the specific enzymes responsible for their glycosylation and further modification were not resolved in this study. Future studies should investigate the enzymatic activities, substrate preferences, and in vivo functions of Cluster-26256.0 and the selected *DFR*-related genes to clarify their contributions to anthocyanin accumulation and petal color formation.

### 4.3. Co-expression network analysis highlights candidate transcriptional regulators of anthocyanin accumulation

The integrated co-expression network provided a correlation-based framework for linking anthocyanin accumulation with structural genes and transcription factors in *Cyclamen* petals. In this study, several glycosylated anthocyanins, including delphinidin-3,5-*O*-diglucoside, cyanidin-3,5-*O*-diglucoside, malvidin-3,5-*O*-diglucoside, peonidin-3,5-*O*-diglucoside, petunidin-3-*O*-glucoside, and petunidin-3-*O*-galactoside, were closely associated with multiple anthocyanin biosynthetic genes in the network. These associations linked the identified anthocyanin derivatives with *CHS*-, *F3H/F3’H*-, *DFR*-, *ANS*-, and *UFGT*-related genes, indicating coordinated variation in metabolite accumulation and transcript abundance across the three materials.

In addition to structural genes, the network contained multiple transcription factor families, including MYB/MYB-related, WRKY, bZIP, AP2/ERF, NAC, TCP, GARP-G2-like, C2C2-Dof, and HB-HD-ZIP members. The presence of these transcription factors indicates that anthocyanin accumulation in *Cyclamen* petals may not be controlled by a single regulatory factor, but rather by a broader transcriptional network. This pattern is consistent with previous studies in ornamental plants, where flower color formation is often associated with coordinated interactions among flavonoid metabolites, biosynthetic genes, and transcriptional regulators. Comparable correlation-based frameworks have been used to prioritize candidate regulators in *Chimonanthus praecox* and *Lilium cernuum* [23,24]. Integrated analyses in Scutellaria baicalensis also identified candidate genes associated with anthocyanin biosynthesis and flower-color variation [25].

Notably, several transcription factor nodes were connected with both key anthocyanin metabolites and structural genes, implying that they may participate in the transcriptional regulation of anthocyanin biosynthesis or modification. However, these associations should be interpreted cautiously because the current network was based on correlation analysis rather than direct regulatory evidence. Therefore, the co-expression network should be regarded as a candidate-screening framework that provides potential regulatory targets for further functional validation, rather than as definitive proof of direct gene regulation.

### 4.4. Candidate MYB Regulators Associated with Anthocyanin Accumulation

Among the transcription factor families identified in the co-expression network, MYB and MYB-related genes deserve further attention because R2R3-MYB transcription factors are widely recognized as important regulators of anthocyanin biosynthesis in many ornamental species [26]. However, the present MYB-annotated transcripts should be regarded as candidate factors until their conserved domains, phylogenetic relationships, and regulatory activities are experimentally verified [27].

In the present study, MYB-Cluster-43585.0 and MYB-Cluster-33560.0 showed close correlation-based associations with anthocyanin metabolites and structural genes. MYB-Cluster-43585.0 showed a progressive increase in expression from BXK to FXK and SHXK, with average FPKM values of 74.15, 161.25, and 197.11, respectively. This expression pattern was directionally consistent with the higher relative accumulation of several glycosylated anthocyanins in colored petals, especially in SHXK. Therefore, MYB-Cluster-43585.0 may be considered a priority MYB-associated candidate for further functional analysis. MYB-Cluster-33560.0 may also be associated with this process, although its specific contribution requires further evaluation.

The potential role of these MYB candidates is supported by studies in other ornamental plants. For example, anthocyanin-associated R2R3-MYB genes have been shown to regulate flower coloration by activating structural genes involved in anthocyanin biosynthesis, either directly or through interaction with bHLH and WD40 partners [28,29]. Based on the observed expression and correlation patterns, MYB-Cluster-43585.0 may be associated with the expression of structural genes involved in late anthocyanin biosynthesis or modification, such as *DFR*- and *ANS*-related genes. Nevertheless, the current evidence is still based on expression patterns and co-expression relationships. Future work should first determine whether MYB-Cluster-43585.0 contains conserved R2 and R3 MYB domains, a bHLH-interaction motif, and subgroup 6 anthocyanin-related activation motifs. Phylogenetic analysis with known anthocyanin-related MYBs, such as *AtMYB75/PAP1*, *AtMYB90/PAP2*, *PhAN2*, *VvMYBA1*, *IsMYBL1*, and *PqMYB113*, would further clarify whether this gene belongs to an anthocyanin-activating MYB clade [30].

To verify the regulatory function of MYB-Cluster-43585.0, additional experimental evidence is required. Yeast one-hybrid, dual-luciferase, EMSA, and transient overexpression assays should be performed to test whether MYB-Cluster-43585.0 directly binds to and activates the promoters of *DFR* or *ANS*-related genes. These experiments will be necessary to determine whether MYB-Cluster-43585.0 functions as an upstream regulator of anthocyanin accumulation or merely represents a co-expressed candidate associated with petal coloration in *Cyclamen*.

## 5. Conclusions

This study integrated anthocyanin profiling and transcriptome analysis to investigate the biochemical and molecular basis of petal color variation in *Cyclamen persicum*. Clear differences in anthocyanin composition and relative accumulation were observed among BXK, FXK, and SHXK. In particular, the dark red petals of SHXK were characterized by the preferential accumulation of multiple terminal glycosylated anthocyanins, indicating that the formation of the dark red phenotype was associated with substantial remodeling of anthocyanin accumulation patterns.

Mapping differentially expressed genes onto the anthocyanin biosynthetic pathway revealed branch-specific transcriptional variation among the three petal-color materials. Among the late biosynthetic genes, the putative anthocyanidin synthase gene Cluster-26256.0 showed an expression pattern directionally consistent with the white, pink, and dark red petal-color classes. Several DFR-related transcripts, including Cluster-46048.3 and Cluster-5522.0, also showed relatively high expression in SHXK. These expression patterns were associated with the higher relative accumulation of multiple terminal anthocyanin derivatives in dark red petals. In addition, the co-expression network highlighted MYB-Cluster-43585.0 and MYB-Cluster-33560.0 as candidate transcription factors associated with anthocyanin accumulation.

Overall, this study provides a metabolite–transcript framework for identifying candidate genes associated with petal color variation in *C. persicum*. The *ANS*-, *DFR*-, and MYB-related transcripts identified here should be regarded as priority candidates for further investigation. Their biochemical functions and regulatory roles require validation through enzyme activity assays, transient expression, and promoter activation analyses.

## 6. Funding

The Key Natural Science Foundation of the Anhui Higher Education Institution of China (Grant No. 2024AH052030) Excellent Young Teacher Training Program of Anhui Province (Grant No. YQYB2024125)

Anhui Provincial Key Project of Humanities and Social Sciences Research in Universities (Grant no.2024AH053489)

Proof of Concept Project at Shanghai Institute of Technology.

## Disclaimer/Publisher’s Note

The statements, opinions and data contained in all publications are solely those of the individual author(s) and contributor(s) and not of MDPI and/or the editor(s). MDPI and/or the editor(s) disclaim responsibility for any injury to people or property resulting from any ideas, methods, instructions or products referred to in the content.

