## Supplementary material for "Integrated anthocyanin profiling and transcriptome analysis identify candidate genes associated with petal color variation in *Cyclamen persicum*": Declaration of competing interests.docx

☒ The authors declare that they have no known competing financial interests or personal relationships that could have appeared to influence the work reported in this paper.

☐The authors declare the following financial interests/personal relationships which may be considered as potential competing interests:
