## Supplementary material for "Integrated anthocyanin profiling and transcriptome analysis identify candidate genes associated with petal color variation in *Cyclamen persicum*": Supplementary material Tables.docx

**Table S1. List of specific primers used for quantitative real-time PCR (qRT-PCR) analysis.**

| Primer name | Upstream sequence(5’-3’) | Downstream sequence(5’-3’) |
| --- | --- | --- |
| *eEF1α* | CTGGTGGTTTTGAGGCTGG | CTGGCCAGGGTGGTTCATGAT |
| Cluster-47815.2 | TGCATATTACTTCCGAGGGGG | CGTACAACACTATTCCATGCTTCC |
| Cluster-30347.7 | CCTCGTGAATCCGATACCCG | AGCAGTTTTTGTGCACATGT |
| Cluster-21594.0 | TGGTAATTTCAAATGTCAAGGTACGA | GAAGCGCCAAATCATGCACA |
| Cluster-19444.10 | GAGCTAGGCTCGTTTCTAGATT | AGTTGAGGTGTCATTGTCTACACT |
| Cluster-25159.7 | GGTGCCTCGGTCAAAAAGGT | GGGACGGGTCATATATGCAGT |
| Cluster-48573.4 | ACTGTTGCTGTGGGAATTGTC | ACACATTCAACGCAACTCACC |
| Cluster-47302.5 | GTTCTCGGGCGTACCTTTGC | AACACACCTTCATAAACCATCAAA |
| Cluster-46532.0 | TGAGAATCAGGTGCTGTGGA | ACTCTCAGTTTGGAGAAAGAACCA |
